# Nanotope: A Graph Neural Network-Based Model for Nanobody paratope prediction

**DOI:** 10.1101/2023.09.27.559671

**Authors:** Xiangpeng Meng, Shangru Li, Rui Li, Bingding Huang, Xin Wang

## Abstract

Nanobodies are artificial antibodies derived from the immune system of camelid species, obtained through artificial processing and isolation of antigen-binding proteins. Their small molecular size and high specificity endow nanobodies with extensive potential applications in various fields. However, determining nanobody paratopes through experimental methods is both costly and time-consuming, while traditional computational approaches often lack sufficient accuracy. To enhance the prediction accuracy, we propose Nanotope, a structurebased model capable of efficiently and accurately predicting paratopes, utilizing graph neural networks and an antibody pretrained language model. Our approach primarily leverages sequence features obtained from the AntiBERTy antibody pretrained language model and three-dimensional spatial structure features of nanobodies as inputs. Employing one-dimensional convolution, EGConv graph convolution, and GATConv convolution, we predict the probability of nanobody paratopes with a much higher accuracy. Our method significantly improves the prediction of nanobody paratopes and requires only nanobody sequences and structural information, without the need for additional antigen information. Source code is freely available at https://github.com/WangLabforComputationalBiology/Nanotope

## I. Introduction

Antibodies are substances known for their high specificity, binding affinity, and capability to neutralize target antigens. Nanobodies, a type of artificial antibodies derived from the immune systems of camelid species through artificial processing and isolation of antigen-binding proteins [1], exhibit comparable binding affinity and specificity to conventional antibodies despite being composed of a single variable domain. However, nanobodies display greater diversity [2] in their complementarity-determining regions compared to conventional antibodies, indicating that determining nanobody- antigen paratopes poses a greater challenge.

Experimental methods [1] for determining nanobody-antigen paratopes are expensive and time-consuming. Alternatively, docking tools [3] offer a faster and less resource-intensive approach, but they often lack accuracy and may not be suitable for high-throughput applications. In recent years, deep learning [4] methods have shown significant advantages in predicting nanobody complementarity-determining regions, particularly when leveraging information from antibody sequences and structures.

Paragraph [1] is a newly developed and freely available tool for antibody epitope prediction based on antibody sequence graphs. It incorporates information about the antibody’s structure by representing it as a graph and using equivariant graph neural network layers to rapidly predict the probability of residues belonging to antigen epitopes. Compared to Parapred [6], a model based on convolutional and recurrent neural networks for antibody paratope prediction, Paragraph demonstrates superior performance.

In this article, we utilize the graph neural network EGConv [8] and a pre-trained language model called AntiBERTy [7], which was trained on 558 million antibody sequences, to construct a structure-based nanobody paratope prediction tool called Nanotope. Nanotope takes the three-dimensional structure of nanobodies as input and uses an efficient graph convolutional neural network to rapidly predict residues belonging to paratope. When compared to existing antibody paratope prediction models, Nanotope outperforms the current state-ofthe-art tool Paragrah.

## II. Methods

### A. Dataset Construction

We collected data from the SabDab [9] antibody structure database, including antigen-antibody complexes and antigen-nanobody complex structures. Each residue in the primary protein was labeled as either a paratope (positive class) or a non- paratope (negative class) based on distances between residues in the complex structure. Complementarity-determining regions on nanobodies and antibodies were defined as residues with distances less than 3.5A° to any heavy atom (excluding hydrogen) on the antigen. Antibodies and nanobodies were renumbered using the IMGT [10] antibody numbering system. Only the variable region Fv sequence and structural information of the heavy chain were retained for antibodies, while nanobodies retained their original sequences. Duplicate sequences were removed, and the data was split into a training set (3:2 ratio for antibody heavy chains to nanobodies) and a testing set consisting of nanobodies.

### B. Graph Construction based on Nanobody Protein Structure

The Cartesian coordinates of *α* carbon atoms for each residue in the nanobody protein structure were extracted from three-dimensional space. Using the K-Nearest Neighbors (KNN) algorithm, we determined the K-nearest residues for each residue based on their Euclidean distances. This information was used to create an adjacency matrix representing each protein structure as a graph, with each amino acid residue as a node. Each node was represented using a 512-dimensional feature vector obtained from the output of the AntiBERTy [7] antibody pre-trained language model. Padding was applied to protein sequences to a length of 140 by adding zero vectors of 512 dimensions to the end of the sequence. The Cartesian coordinates of the padded nodes were set to (0, 0, 0) in threedimensional space.

### C. Graph Neural Network Model

The constructed nanobody graphs served as inputs to the model. The initial 512-dimensional vectors obtained from AntiBERTy [7] were processed through a three-layer onedimensional dilated convolutional structure, with each layer followed by one-dimensional batch normalization. Subsequently, a three-layer efficient graph convolutional layer and one Graph Attention convolutional layer [11] were applied. The efficient graph convolutional layer consisted of an EGConv layer [9], one-dimensional batch normalization, and ReLU activation. Finally, a four-layer feed-forward network (FFN) with ELU activation was used for classification. The overall structure of the model is shown in Figure 1.

**Fig. 1:**
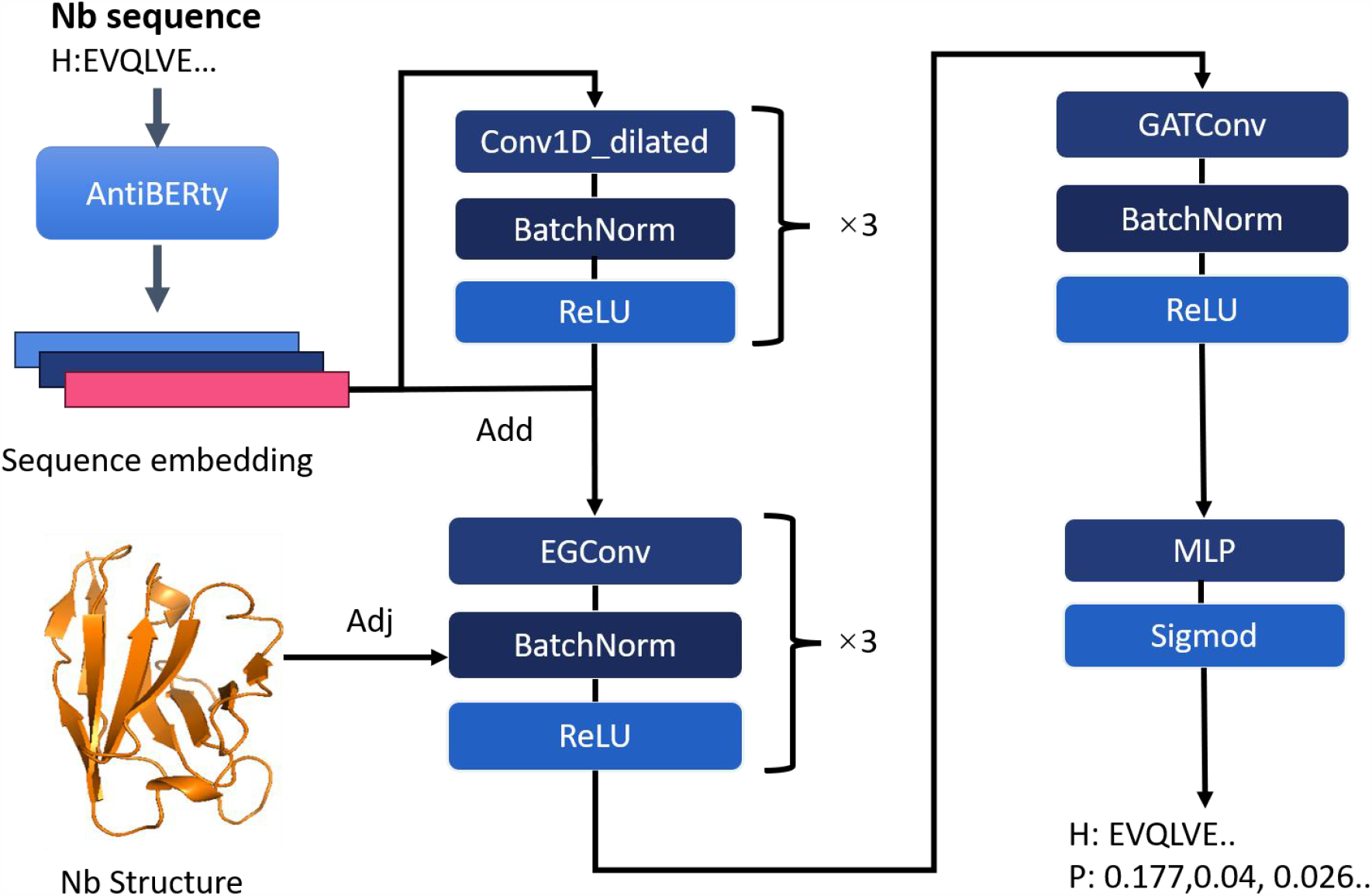
The Model Architecture of Nanotope. The antibody sequences are transformed into contextual embeddings using the pretrained language model, AntiBERTy. Then, the nanobody structures are converted into graphs using the K-Nearest Neighbors (KNN) algorithm. As shown in the figure, Nanotope utilizes graph neural networks to predict the probability of each residue being a paratope.

### D. Model Training

To address the imbalanced binary classification task, we used the binary cross-entropy loss function during model training. The hyperparameters were set as follows: the onedimensional dilated convolution had kernel sizes [5, 5, 3] and a dilation rate of 4. The EGConv [8] had 8 heads and 8 B basis weights, using 5 aggregators [‘mean’, ‘sum’, ‘std’, ‘max’, ‘min’]. The GATConv [11] had 2 heads. The model was trained for 10 epochs with a batch size of 64. The number of nearest neighbor edges was set to 16. The Adam optimizer was used to optimize model parameters, and a cosine annealing learning rate scheduler [12] was employed with an initial learning rate of 0.001.

## III. Results

To ensure the fairness of evaluation, this model was tested using the nanobody data that was not used during the training process. We employed a 10-fold cross-validation [13]technique to assess the performance of our model. To ensure that the validation sets used in cross-validation were exclusively composed of nanobody data, both the nanobody data and heavychain antibody data were randomly divided into 10 sets. The model was then trained ten times, each time using a different nanobody dataset as the validation set and the remaining nine sets of nanobody and heavy-chain antibody data as thetraining set. The results of the cross-validation are presented in the supplementary materials.

Since there are currently no specialized methods for predicting nanobody paratope, our study compared our model with other representative antigen-antibody paratope prediction methods, including Paragraph [5], Parapred [7], ProBERT [14], and our constructed Baseline method. Baseline method involves calculating the frequency of nanobody training set residues that are positive instances according to IMGT [10] numbering and using this frequency to replace the probability of the same encoded positions and residues in the test set. Figure 2. displays the AUC and PR AUC curves for all methods except Baseline.

**Fig. 2:**
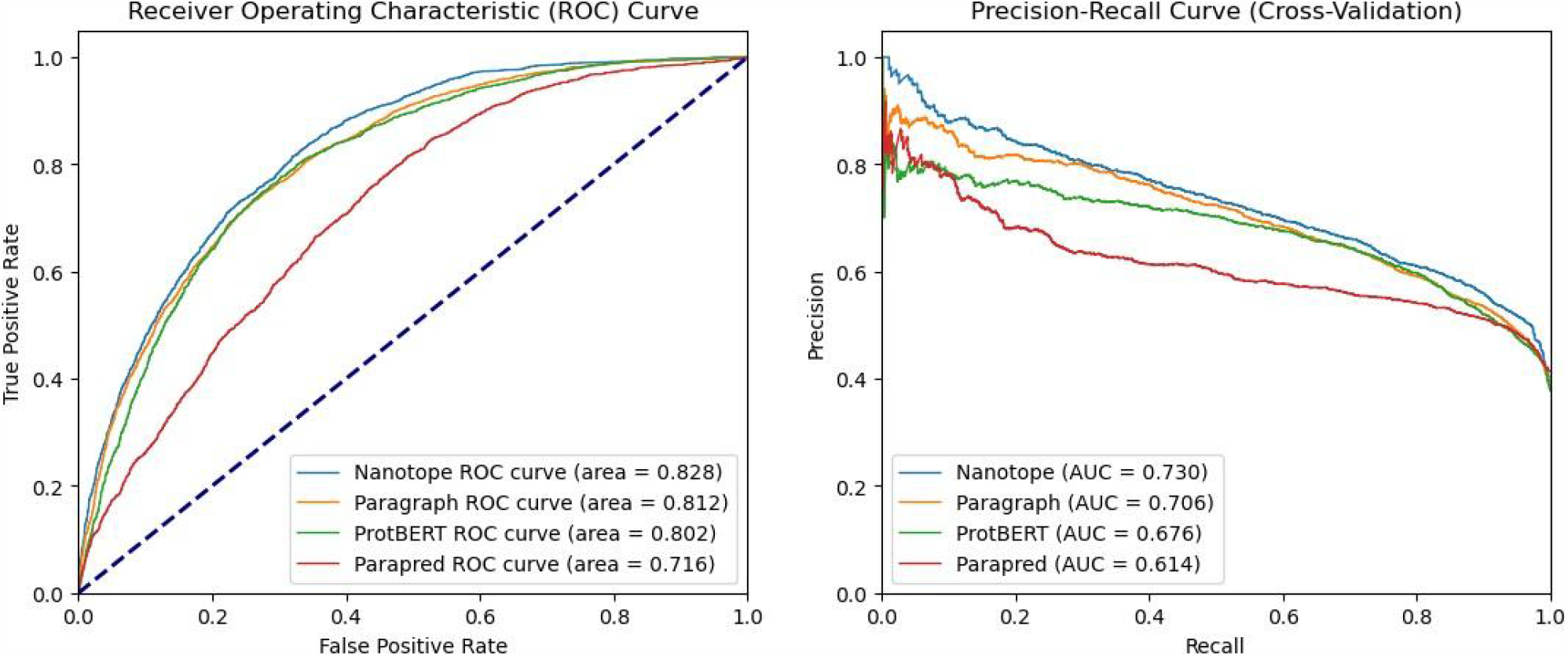
Nanotope outperforms publicly available tools for paratope prediction. **(A)** receiver operating characteristic (ROC) curves and **(B)** Precision-recall for paratope prediction by Paragraph [5], Parapred [6], ProtBERT [14] and Nanotope.

Table 1 presents the comparison results, indicating that our model outperforms existing antibody paratope prediction methods and also surpasses the basic Baseline method. The superiority of our model in predicting nanobody paratope highlights its advantage in this particular task. All protein data bank (PDB) codes can be found at ./tree/main/Data.

**TABLE I:** Comparison of paratope prediction methods.

| Method | ROC AUC | PR AUC |
| --- | --- | --- |
| Baseline | 0.795 | 0.674 |
| Parapred | 0.716 | 0.614 |
| ProBERT | 0.802 | 0.676 |
| Paragraph | 0.811 | 0.705 |
| **Nanotope** | **0.827** | **0.730** |

Note: All evaluations are conducted on the CDR region of the nanobody model structures. All results are derived from the nanobody dataset that we processed. As there are currently no existing models for predicting nanobody, all the models used in the evaluation are assessed using their respective trained antibody prediction models. Specifically, the ProBERT [14] model’s results are obtained through fine-tuning on the nanobody dataset.

## IV. Discussion

We conducted an analysis of the final layer’s Graph Attention Network [12] convolutional layer in our model. The main approach involved extracting the attention weights of the edges connected to the GAT layer’s nodes and visualizing them (Figure 3). We noticed that although the edges in our model were defined as the K nearest neighbors, we did not explicitly input the ranking information of the nearest edges. Instead, we only considered the K nearest edges as the edges of the nodes in the graph.

**Fig. 3A.**
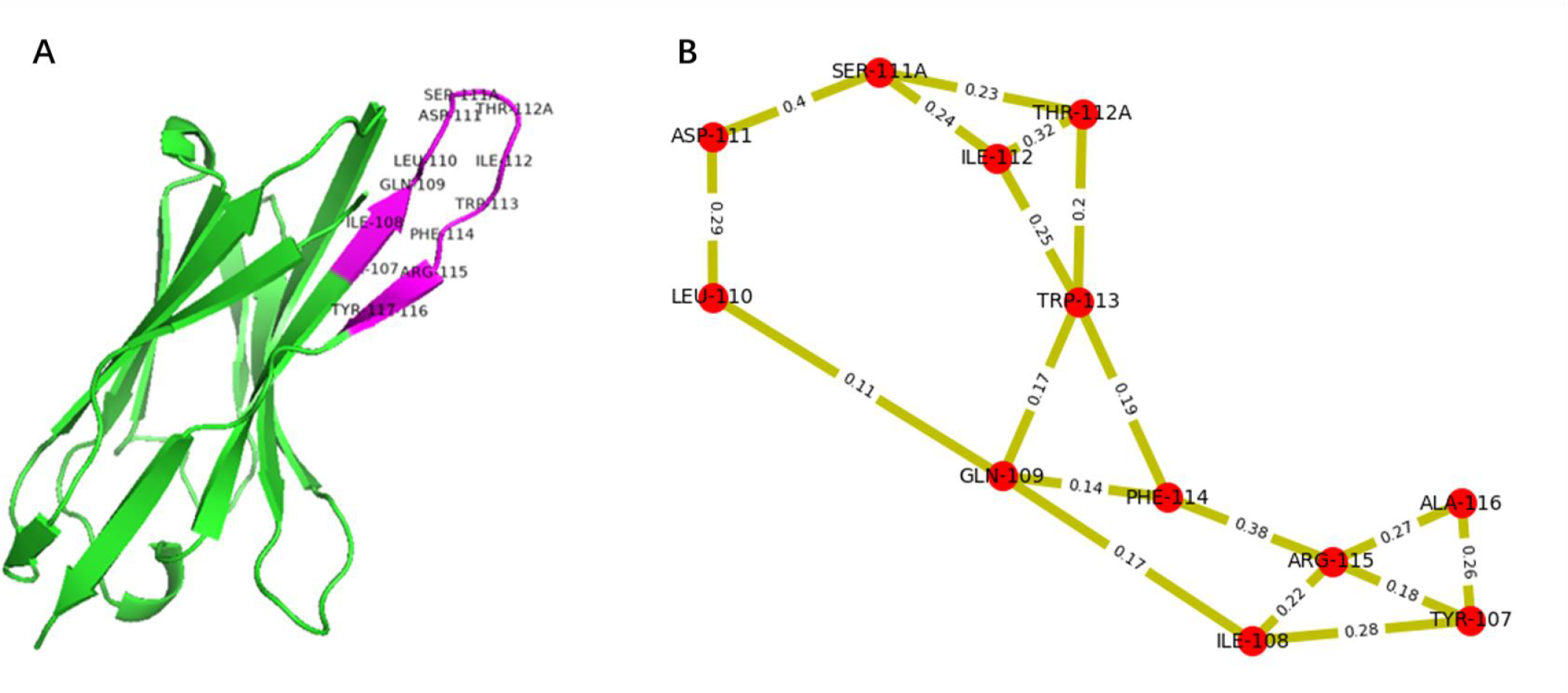
shows the crystal structure of a nanobody, where the purple region represents the CDR regions (PDB: 6h72). Fig. 3B shows the visualization result of the attention part (CDR3) of the Nanotope model. The attention results can learn the spatial representation of the nanobody structure. Taking the node “ARG-115” in the CDR3 region as an example, the model pays more attention to the two residues “PHE-114” and “ALA-116”, which are closest to this node, with attention coefficients of 0.38 and 0.27, respectively. On the other hand, the model assigns less attention to the two residues “ILE-108” and “TYR-107”, which are relatively distant from this node, with attention coefficients of 0.22 and 0.18, respectively.

For instance, concerning node APG-115 (Figure 3), the model assigned higher attention to certain edges, which were later confirmed to connect nodes closer to APG-115. Thus, for our model, edges with larger attention coefficients tend to connect two more adjacent nodes, indicating that the model learned the spatial structure of the data. This finding suggests that even without explicitly providing the spatial structure of the nanobodies, the model is capable of discerning spatial distances and using them as evaluation criteria.

In the more challenging task of nanobody paratope prediction, traditional antibody paratope prediction models exhibit limited performance. To address this, we proposed a deep learning model for predicting nanobody-antigen paratopes. Our results demonstrate that the model successfully captures the structural characteristics of nanobodies and achieves state-of-the-art performance in the task of nanobody epitope prediction. The application of deep learning techniques has proven effective in improving the accuracy and efficiency of nanobody paratope prediction, showcasing the potential for further advancements in this field.

